# APEX-Gold: A genetically-encoded particulate marker for robust 3D electron microscopy

**DOI:** 10.1101/2020.10.06.328831

**Authors:** James Rae, Charles Ferguson, Nicholas Ariotti, Richard I. Webb, Han-Hao Cheng, James L. Mead, Jamie Riches, Dominic J.B. Hunter, Nick Martel, Joanne Baltos, Arthur Christopoulos, Nicole S. Bryce, Maria Lastra Cagigas, Sachini Fonseka, Edna C. Hardeman, Peter W. Gunning, Yann Gambin, Thomas Hall, Robert G. Parton

**Author notes:** Author for Correspondence: Robert G. Parton, Institute for Molecular Bioscience, The University of Queensland, St Lucia, Queensland 4072, Australia. These authors contributed equally (James Rae, Charles Ferguson).

## Abstract

Genetic tags allow rapid localization of tagged proteins in cells and tissues. APEX, an ascorbate peroxidase, has proven to be one of the most versatile and robust genetic tags for ultrastructural localization by electron microscopy. Here we describe a simple method, APEX-Gold, which converts the diffuse oxidized diaminobenzidine reaction product of APEX into a silver/gold particle akin to that used for immunogold labelling. The method increases the signal to noise ratio for EM detection, providing unambiguous detection of the tagged protein, and creates a readily quantifiable particulate signal. We demonstrate the wide applicability of this method for detection of membrane proteins, cytoplasmic proteins and cytoskeletal proteins. The method can be combined with different electron microscopic techniques including fast freezing and freeze substitution, focussed ion beam scanning electron microscopy, and electron tomography. The method allows detection of endogenously expressed proteins in genome-edited cells. We make use of a cell-free expression system to generate membrane particles with a defined quantum of an APEX-fusion protein. These particles can be added to cells to provide an internal standard for estimating absolute density of expressed APEX-fusion proteins.

## Introduction

Genetic tags for electron microscopy (EM) have made protein ultrastructural localization possible within the three-dimensional (3D) environment of cells, tissues, and whole organisms (Ariotti et al., 2015; Han et al., 2019; Lam et al., 2015; Martell et al., 2017; Martell et al., 2012; Tsang et al., 2018). APEX, a modified ascorbate peroxidase derived from soybean, is a simple to use highly versatile marker for EM detection. The wide applicability of its use is demonstrated by studies utilizing APEX and its derivative, APEX2, for detection of fusion proteins in cells, *Drosophila*, zebrafish, and mice (Ariotti et al., 2015; Hirabayashi et al., 2018; Li et al., 2020; Meiring et al., 2019). APEX has also been combined with nanobody-based detection systems for rapid localization of GFP- and mCherry-tagged proteins (Ariotti et al., 2015; Ariotti et al., 2018), and used to detect protein interactions using split GFP and nanobodies or using a novel split-APEX system (Ariotti et al., 2018; Han et al., 2019). These applications are compatible with 3D EM techniques in which the reaction product can be produced within the depth of the specimen, rather than on the surface as occurs with labeling on sections (Martell et al., 2017; Martell et al., 2012), and can also be used with methods that require cryo-preservation (Tsang et al., 2018). Despite its versatility, the relatively diffuse DAB reaction product can make distinguishing the APEX reaction from electron dense cellular components difficult and often requires an expert to interpret the images. This is especially the case when APEX-tagged proteins have multiple subcellular localizations, for example, a soluble and a membrane-localised distribution (Follett et al., 2016). In addition, a method which allows researchers to obtain a particulate signal from a genetic tag that could be equated to the actual number of antigens present would be a huge breakthrough in the field.

In this study, we describe the use of a method for converting the DAB reaction produced by a cytoplasmically exposed APEX tag into a particulate marker. This method produces an easily detectable gold particle at the site of the fusion protein with high resolution and specificity, allowing simple correlative light and electron microscopy (CLEM) and use in 3D EM techniques. In combination with *in vitro* synthesized APEX-fusion protein particles this modification allows estimation of the density of a protein *in situ* in cells and tissues.

### Design

In order to convert the DAB reaction product to a particulate marker, we tested a number of protocols with a particular focus on silver/gold (Ag/Au) enhancement methods originally developed for amplification of the DAB signal obtained with peroxidase-labeled antibodies on histological sections. The criteria for enhancement of the APEX-DAB reaction product for EM were; 1) production of a uniformly-sized particle with high specificity, 2) high sensitivity with low background and no self-nucleation, 3) high resolution, 4) ease of use, i.e. using conventional fixation and processing schemes, readily available laboratory reagents, and in ambient light conditions rather than a darkroom. The optimal protocol which satisfied these criteria was a modified silver/gold enhancement method (Sedmak et al., 2009) as shown schematically in Fig. 1A, similar to that used to visualise luminal APEX (Mavlyutov et al., 2017). After fixation and a conventional incubation with DAB/H2O2 to reveal the oxidized DAB reaction product, cells were incubated sequentially with a silver nitrate solution (containing hexamethylenetetramine and disodium tetraborate) and then with gold chloride. This resulted in local production of stabilised gold particles in the range of 10-15 nm in diameter as the argyrophilic oxidized DAB reaction product converts the metal salts to colloidal particles at the site of the fusion protein. Gum arabic was included to provide consistent uniform nucleation.

**Figure 1.**
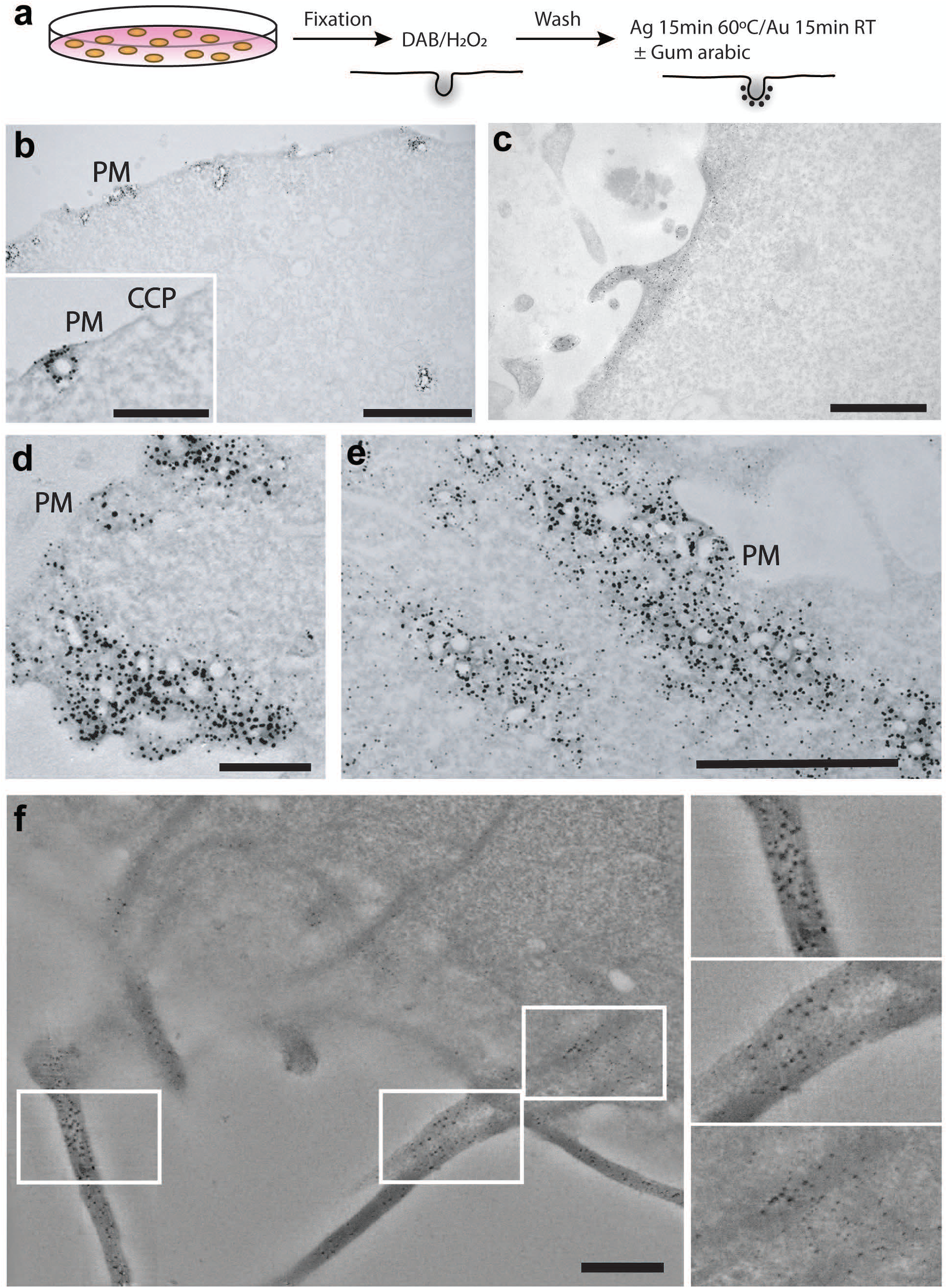
**A;** Schematic of the APEX-Gold method. Cells were transfected with Cavin4-APEX2 (B, D,E) (control light microscopy experiments are described in Supplementary Fig. 4) or LifeAct-APEX2 (C,F), fixed, treated with DAB and then incubated with Ag/Au reagents in the presence of gum arabic. **B,D,E;** Low (B) and higher (D,E, inset in B) magnification views of caveolae labeling. C; Labelled actin filaments. F; Optical slices through tomogram of LifeAct-APEX2 expressing cells. APEX-Gold particulate reaction product can be observed tightly associated with and throughout the actin bundles in 3-dimensions. Note the uniform gold label, the lack of background, and high signal-to-noise. PM, plasma mem brane; CCP, clathrin-coated pit. Bars, B, 2µm, B inset 500 nm; C, 1 µm; D, 500 nm; E, 1 µm; F, 500 nm.

## Results

We applied this localization method to Cavin4-APEX2 which is associated with cell surface caveolae (Fig. 1B,D,E and Supplementary Fig. 1A,B), to LifeAct-APEX2 that allows ultrastructural detection of actin filaments (Fig. 1C and Supplementary Fig. 1C,D), and to A1AR-APEX2 a G-protein-coupled receptor (Supplementary Fig. 6D). The APEX-Gold method satisfies the criteria of specificity, low background, sensitivity, resolution, and ease of use. Untransfected cells show negligible silver/gold particles (Supplementary Fig. 6A) and labeling is tightly restricted to caveolae with an average of 18.4 nm from the caveolar membrane to the centre of the particulate reaction product (Supplementary Fig. 5A,B). The experimental process is very simple and robust; all incubations are done in the light and require no specialist chemicals or equipment. The method has been used successfully in over 20 different biological replicate experiments using the Cavin4-APEX2 and LifeAct-APEX2 systems. Critically, it results in an unambiguous particulate reaction product that is clearly definable without expert interpretation.

The clarity and density of the APEX-Gold enhanced particulate signal makes the method compatible with 3D EM methods including 3D electron tomography (Fig. 1F) array tomography scanning EM (Supplementary Fig. 2A,B), and focussed ion beam (FIB) EM (Supplementary Fig. 2C). The gold particles can also be resolved by automated segmentation (WEKA Image J/FIJI plug-in, Supplementary Fig. 2 C’,D’). The APEX-Gold method is also simple to use in CLEM approaches (Fig. 2A-D) and can be checked by dot blot in parallel to the EM experiment to ensure the protocol is standardized (Supplementary Fig. 7B; see Materials and Methods for a standard protocol). We also tested the compatibility of the method with freeze substitution/low temperature embedding (Supplementary Fig. 6B,C). This makes APEX-Gold potentially compatible with double labeling, with the APEX fusion protein being expressed endogenously in the cells and then sections labeled for other proteins of interest.

**Figure 2.**
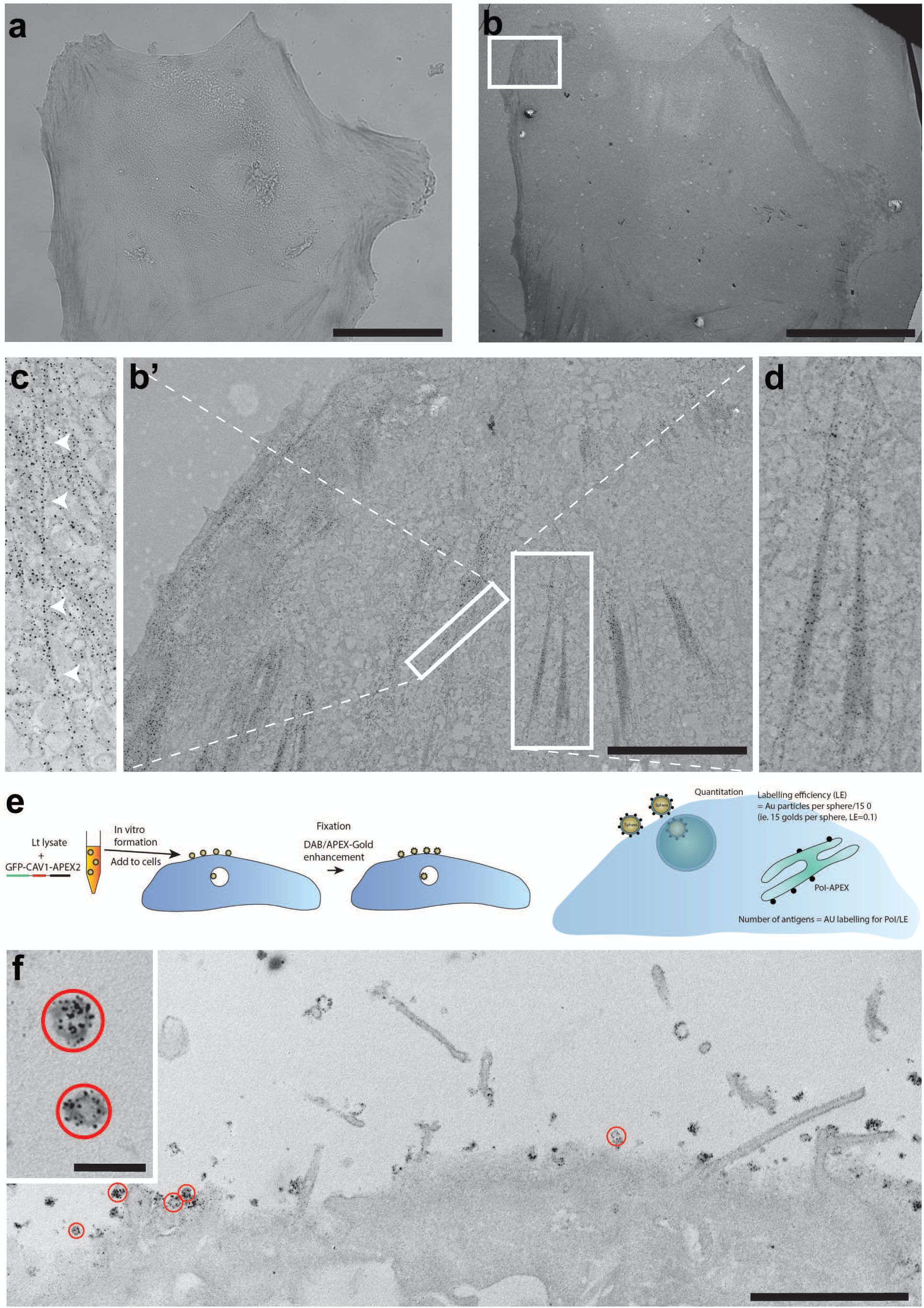
**A;** Light microscopic detection of Tpm3.1-APEX2 after APEX-Gold DAB/Ag/Au detection. **B;** Low magnification EM showing a basal section of the same cell. **B’,D;** Higher magnification views of gold labeled stress fibers from boxed region. **B’,C;** Gold labeling follows individual actin filaments from boxed region. E; Schematic explaining the cell-free caveolae-APEX2-Gold system. F; A431 cells were incubated with *in vitro* synthesized CAV1-APEX2 cell-free caveolae for 5 min at 37°C before fixation and processing for APEX-Gold detection. Note the Ag/Au labeling of the surface-associated cell-free caveolae circled in red. Bars, A,B, 50 µm; B’, 5 µm; F, 2 µm, F inset 200 nm.

Next we investigated the suitability of the APEX-Gold method for the detection of proteins expressed at levels close to, or even at tracer levels below endogenous expression levels. For this we utilized a recently characterized CRISPR generated mouse line that expresses tropomyosin 3.1 (Tpm3.1) C-terminally tagged with APEX2 from the endogenous locus (Meiring et al., 2019). In these mice, exon 9d of the tropomyosin gene is fused to APEX2 via a 10 amino acid linker ((Meiring et al., 2019); Supplementary Fig. 3A). Western blotting of mouse embryonic fibroblasts (MEFs) prepared from Tpm3.1–APEX2 heterozygous mice showed that the levels of the fusion protein is approximately 5% of the levels of the endogenous Tpm3.1 protein (Supplementary Fig. 3B,C). We then subjected the MEFs to the APEX-Gold method for detection of the fusion protein. Light microscopy studies revealed that specific labeling was associated with putative stress fibers (Fig. 2A) and this was confirmed by correlative EM (Fig. 2B-D) that showed a dense and highly specific particulate labeling associated with bundles of actin, with an average of 10.6 nm from the particle to the centre of bundle mass (Supplementary Fig. 5C-E). In addition, particulate labeling followed distinct tracks suggestive of individual actin filaments (Fig. 2C and Supplementary Fig. 3D-I). These results clearly demonstrate the power of the APEX-Gold technique for detecting proteins at low expression levels and/or proteins expressed from endogenous loci.

Finally, we investigated whether we could develop a system to act as an internal control for the APEX-Gold method and to allow quantitative comparison with cellular APEX-tagged proteins of interest. We made use of the ability of mammalian caveolin-1 (CAV1), the major structural protein of caveolae, to generate nanovesicles with a defined number of caveolin proteins when expressed in a cell-free system (Jung et al., 2018)(see scheme in Fig. 2E). CAV1-APEX2 was expressed in a cell-free *Leishmania* lysate (GFP-tagged and untagged; Supplementary Fig. 7C) and the GFP-tagged protein characterized by fluorescence correlation spectroscopy (FCS) (Supplementary Fig. 7A). The resulting particles contained a quantum of fluorescence consistent with approximately 150 GFP-CAV1-APEX2 molecules per vesicle as observed in previous studies with GFP-CAV1 (Jung et al., 2018). We then compared the enzymatic activity of the cell-free synthesized GFP-CAV1-APEX fusion protein with commercial horseradish peroxidase. The cell-free synthesized APEX2 fusion protein had higher activity per µg of protein than the commercial HRP preparation (Supplementary Fig. 7B). Enhancement of the signal using the APEX-Gold protocol caused a slight increase in sensitivity of detection and a colour change in the dot blot. We then investigated the use of the uniformly sized *in vitro* generated GFP-CAV1-APEX2 particles for comparative studies in cells. The lysate containing the GFP-CAV1-APEX2 particles were added to cultured A431 cells for 5 minutes at 37°C prior to fixation, DAB treatment, and APEX-Gold enhancement. As shown in Figure 2F the GFP-CAV1-APEX2 particles were observed on the surface of the cells and in endosomes and are clearly decorated with the APEX-Gold enhanced particles. This allows direct readout of antigen density by comparison to known standards. While we demonstrated this principle by using exogenously added Caveolin-APEX vesicles, an APEX-tagged antigen generated by CRISPR technology that is present at a known density in a specific region of the cell could be also used as a standard.

## Discussion

Here we present a simple technique for identification of a protein of interest by transmission EM using a genetic tag to generate a particulate signal. The method has high resolution, low background, and is sensitive enough to detect proteins expressed at or below endogenous levels. We provide a defined standard for calibration to ensure reproducibility, and show that the method is compatible with 3D EM techniques such as serial blockface SEM, array tomography, focussed ion beam SEM, and electron tomography.

The APEX system has proven to be a powerful new addition to the biologist’s arsenal of techniques with wide applications in EM and in proteomics. The original APEX system involving a fusion with the protein of interest has been extended to new modifications including nanobody-based detection of fluorescent proteins and protein complexes and a split-APEX which only yields an active enzyme if brought together by two partner proteins. All of these applications are compatible with the APEX-Gold method for conversion of DAB to an Ag/Au particle. The APEX-Gold method is so simple that it should be used in parallel to light microscopy as an initial step in protein localization. In fact, through the use of nanobody-APEX constructs a co-transfection with a GFP-tagged protein can yield correlative light and EM images in a very short timeframe. The technique produces a particulate signal which is clearly distinguished from any cellular feature and is readily quantitated. By combining with an APEX-fusion protein of known signal density (here demonstrated by cell-free generated particles of a defined APEX density as a calibration standard) it is possible to quantitatively compare particle density to antigen density. While we demonstrated this principle by using exogenously added cell-free synthesized caveolin-AP vesicles, an APEX-tagged antigen generated by CRISPR technology that is present at known density in a specific region of the cell could also be used as a standard.

We were able to use the APEX-Gold method to localize low levels of APEX2-tagged proteins and to estimate the sensitivity of the technique. We utilized cells from mice genetically modified to express APEX2 fused to tropomyosin 3.1 which have been previously characterised in detail (Meiring et al., 2019). Embryonic fibroblasts from the mice express the Tpm3.1-APEX2 fusion protein at approximately 5% of endogenous levels. Tpm3.1 is expressed at a concentration of 30µM (Meiring et al., 2018) and so 5% represents a concentration of 1.5µM. This equates to a cellular Tpm3.1-APEX2 copy number of approximately 1 million Tpm3.1 homodimers per cell with 250,000 predicted to be associated with actin and the remainder cytosolic (Meiring et al., 2018). This level of expression resulted in an excellent signal to noise ratio for the APEX-Gold particles on actin filaments. This demonstrates the remarkable efficiency of antigen detection and lays the foundation for truly quantitative immunoEM in cells and tissues for the first time.

APEX-Gold represents a significant advance in protein distribution analysis by EM. Compared with current APEX methods which rely on subjective interpretation of a diffuse DAB reaction product, the improved signal-to-noise ratio and a particulate, quantitative readout, indicate that this method should become the gold-standard for localization of genetically tagged proteins in electron microscopy.

## Limitations

The method used here, which we have optimized for APEX-localization, is based on well-established methods for histological and EM visualization of the insoluble diaminobenzidine product of peroxidase (Adams, 1981; Danscher and Norgaard, 1983; Dobo et al., 2011; Newman and Jasani, 1998; Pohl and Stierhof, 1998), combined with the methods of silver reduction and gold toning developed for photography in the 19^th^ century (for review see (Ellis, 1975)). These methods have been developed and used in many laboratories to allow enhancement of the DAB reaction for light microscopic applications, for converting the DAB signal to an easily detected particulate marker to allow discrimination from any cellular structures, and to increase the sensitivity of DAB detection (Dobo et al., 2011; Newman et al., 1983; Sedmak et al., 2009). The 3 principle steps involve 1) production of the polymerized DAB product, 2) the reduction of silver ions to submicroscopic metallic silver by the argyrophilic DAB polymer, and 3) substitution of metallic silver by gold (gold toning), through the simple reaction Ag + Au^+^ -> Ag^+^ + Au to produce the more inert gold particle that can resist subsequent osmium treatment. A further refinement is the use of gum arabic which was introduced for gold enhancement by (Danscher, 1981; Danscher and Norgaard, 1983) to allow precise control of silver/gold nucleation and more uniform particle generation.

While the benefits of the APEX-Gold method are clear, this approach still comes with certain caveats that must be considered prior to performing these analyses. First, it is necessary to take care to precisely control the development time for particulate deposition with the silver enhancement and subsequent gold toning. We optimized the development conditions to suit the size distribution of the particulate reaction product for detection of APEX-tags at a local density that was suitable for caveolar-localized proteins, tropomyosin decoration of actin filaments and actin bundles. Increased development times can facilitate visualization of the signal but can cause appearance of artefactual self-nucleated particles at sites lacking APEX (hence the need for careful control experiments) and can cause more variable, and fused, gold particle production. Second, despite carefully controlling development times, there still remains some variability in particle size (see for example Supplementary Fig. 1A,B). This can cause potential difficulties for quantitation and for applications in double labeling, for example when the APEX-Gold method is combined with on-section labeling of Tokuyasu sections or freeze-substituted Lowicryl sections. This variability can come from two sources. Firstly, there is some inherent variability in the nucleation giving rise to variation in particle size. A second source of variation may be specific to the system being studied. Careful examination of the labeling pattern for cavin4 reveals larger particles close to the caveolar membrane, and smaller particles in the neighbouring cytosol. This difference in size may be explained by the existence of cavin4 in different oligomeric states on the caveolar membrane and in the cytosol (Kovtun et al., 2014), ie, it reflects a real biological difference which could be used to infer the state of the protein in particular sites. This problem can potentially be avoided by expressing tracer amounts of APEX-tagged proteins, as shown here for cells expressing an APEX-tagged Tropomyosin isoform at tracer levels below endogenous levels. Despite these caveats, the APEX-Gold method can provide superior signal-to-noise ratios, allow absolute quantitation of particulate signals, is compatible with 3D EM methods and automated segmentation, and has resolution equal to, or higher, than immunogold labeling.

## Acknowledgments

The authors acknowledge the help of staff and use of facilities in the Microscopy Australia NCRIS Facility at the Centre for Microscopy and Microanalysis at The University of Queensland. The authors acknowledge the use of the Cryo Electron Microscopy Facility through the Victor Chang Innovation Centre, funded by the NSW government, the Electron Microscope Unit within the Mark Wainwright Analytical Centre at UNSW Sydney and Microscopy Australia. This work was supported by the National Health and Medical Research Council of Australia (grants APP1140064 and APP1150083 and fellowship APP1156489 to R.G.P.; APP1185021 to N.A.). RGP is supported by the Australian Research Council (ARC) Centre of Excellence in Convergent Bio-Nano Science and Technology. This work was also supported by an Australian Department of Industry, Innovation and Science Cooperative Research Centre Project (CRC-P) grant to P.W.G. and E.C.H. and grants from the Australian Research Council (ARC grant DP160101623), the Australian National Health and Medical Research Council (NHMRC grant APP1100202, APP1079866), and The Kid’s Cancer Project to P.W.G. and E.C.H.

## Author Contributions

R.G.P. designed the study. J.Rae and C.F. optimised silver enhancement methods and carried out electron microscopy experiments with assistance from R.W. (freeze substitution), J.Riches and N.A. (electron tomography). D.H. performed the cell-free synthesis experiments, guided by Y.G., N.M. and T.H. designed constructs and performed the cloning. J.B., guided by A.C., performed optimisation experiments with APEX-tagged membrane proteins. J.Rae and T.H. performed light microscopy experiments. P.G., M.L.C., N.S.B., and E.C.H. generated the Tm3.1-APEX2 mouse. M.L.C. and N.S.B isolated and characterized the Tm3.1-APEX2 expressing mouse embryonic fibroblasts. R.G.P., J.Rae, and N.A. analysed the data, collated figures, and wrote the manuscript with input from all the other authors.

## Competing Financial Interests

The authors declare no competing financial interests.

## Methods

### Cell Culture

BHK (ATCC), A431 (ATCC), MEF (as described in (Meiring et al., 2019)) and Flp-In-CHO (Invitrogen) cells were maintained in Dulbecco’s Modified Eagle Medium (DMEM), supplemented with L-glutamine, and 10% fetal bovine serum (FBS) at 37°C with 5% CO2. CHO cells stably expressing the human adenosine A1 G protein-coupled receptor (A1AR) with C-terminally labeled APEX2 were generated as described previously (Baltos et al., 2016). Expression was maintained by addition of 500 μg/mL hygromycin-B to culture medium.

### Plasmid Construction

Cav1-APEX2, Cav1-EGFP-APEX2, Cavin4-APEX2-P2A-mKate2 and LifeAct-APEX2-P2A-mKate2 were produced using the multisite gateway system (Invitrogen) by recombination of pME-Cavin4, pME-Cav1, pME-LifeAct, p3E-APEX2-P2A-mKate2, p3E-APEX2 and pCSDEST2. Full details and unique repository identifiers are given in STAR methods below. A1AR-APEX2 was produced by gateway cloning, whereby human A1AR in pENTR/D/TOPO vector was recombined into pEF5/FRT/V5-DEST-APEX2. The resultant expression vector encoded the human A1AR with C-terminally tagged APEX2, adjoined by a glycine-serine rich linker. GFP-CAV1 was produced by cloning into pCellFree_G03 (Gagoski et al., 2015) from human ORFeome library described in (Skalamera et al., 2011).

### Transfection

Cells were seeded into 35mm tissue culture dishes ON, then transfected with Lipofectamine 3000 as per manufacturer’s instructions. Cells were left for 24 hours then fixed and processed for EM 24 hours later.

### Electron Microscopy

#### eLine

SEM array tomography was carried out in an Electron Beam Lithography system, the Raith eLINE PLUS. The system is equipped the dual detector (Inlens, SE) and a laser interferometric stage. Prior to the image acquisition the scan field was calibrated at 25,000x with the laser stage. The displayed image was captured at 2 kV with 30 µm aperture (beam current of 30 pA), using the SE detector, at a working distance of 2.2 mm. The use of laser interferometric stages allows near perfect stitching of scan fields by overlapping just 5 pixels at the edge.

#### FIB

FIB-SEM tomography of the sample was carried out on a FEI Scios DualBeam FIB-SEM system equipped with a 30 kV Ga+ column and a Pt gas injection system (FEI). The sample was tilted to 52° so that its top surface was aligned parallel to the focal plane of the FIB. A protective layer of Pt was deposited onto the top surface directly above the volume of interest. A trench was milled at one end of the volume of interest using a 7 nA beam current. The purpose the trench was to provide the SEM with an unobstructed view of the exposed cross-section and to allow for the escape of sputtered material. Fiducial marks were milled into both the top surface of the sample and the sidewall of the trench. The surface of the exposed face was planarized by milling using successively lower ion beam currents, down to 100 pA. The Auto Slice and View automation package (FEI) was used to sequentially mill away 10 nm thick segments of material followed by SEM imaging of each newly exposed surface using the In-lens Trinity ‘T1’ BSE detector. An electron beam acceleration voltage of 2 kV and current of 50 pA was used. The stage remained stationary during the entire sequence so that the surface of each cross-section is tilted 38° from the SEM column and in-column detector. The respective fiducial marks were detected by the automation package prior to each milling and imaging step and used to correct for image drift of the electron and ion beam, respectively.

#### Tomography

Thick 200 nm sections were cut on a Leica UC6 ultramicrotome. Grids were assembled into an Autogrid (Thermo Fisher) and loaded onto a 200 kV Thermo Fisher Talos Arctica fitted with a Falcon 3EC (Thermo Fisher) camera operated in linear mode and at room temperature. Bidirectional dual axis tilt series were acquired at 1° increments from -60° to +60 ° under the control of Tomography software (Thermo Fisher). Tilt series were reconstructed using weighted-back projection with IMOD.

### DAB treatment and APEX-Gold enhancement

DAB treatment and APEX-Gold enhancement were performed using a modification of the method of Sedmak et al, originally developed to enhance immunoperoxidase staining of tissues (Sedmak et al., 2009). Cells grown in 3 cm dishes were fixed with glutaraldehyde (2.5%) in PBS, then washed in PBS and then in cacodylate buffer, pH 7.35. Fixed cells were incubated in freshly prepared 0.05% DAB solution in cacodylate buffer for 10 min at RT followed by incubation with 0.05% DAB solution containing 0.01% H2O2 for 30 min at RT. The cells were then washed with cacodylate buffer and further fixed in 2.5% GA in cacodylate buffer for 1 hour at 4°C to stabilize the DAB reaction product. After washing with cacodylate buffer, cells were immediately processed for APEX-Gold enhancement.

Cells in dishes were washed in triple distilled water (H2O) for 4 x 15 min to remove phosphates and reduce artefactual silver/gold particle deposition. Cells were then blocked prior to silver enhancement with an aqueous solution containing 1% BSA and 20 mM Glycine for 20 min. Dishes containing cells were prewarmed at 60°C for 10 min. An enhancement solution containing 3% hexamethylenetetramine (C6H12N4) in H2O, 5% silver nitrate (Ag NO3) in H2O, and 2.5% disodium tetraborate (Na2B4O710H2O) in H2O, mixed in a ratio of 20:1:2 was prewarmed and added to the cells and incubated for 15 min at 60°C. After washing in H2O (3 x 5 min) cells were then incubated with 0.05% tetrachlorogold (III) acid trihydrate (AuHCl43H2O) in H2O for 5 min at RT, washed in H2O, and incubated with 2.5% sodium thiosulphate for 4 min at RT. In some experiments, as indicated, the enhancement solution was mixed at a 1:1 ratio with an aqueous 50% gum arabic solution. These dishes were later rinsed with warmed H2O to facilitate removal of residual gum arabic.

Cells were then postfixed with 1% osmium for 2 min, then serial dehydrated with increasing percentages of ethanol. Cells then underwent serial infiltration with LX112 resin in a Pelco Biowave then incubated at 60°C for 24 hours. Ultrathin sections were attained on a ultramicrotome (UC6, Leica), and imaged using a JEOL1011 transmission electron microscope at 80 kV.

### Freeze substitution

Cells grown on Thermanox coverslips were DAB treated and underwent the APEX-Gold enhancement described earlier. Cells were then infiltrated with 2.1 M sucrose at 4°C ON, fast frozen in liquid nitrogen and freeze substituted in 0.2% uranyl acetate in methanol, then washed in methanol and infiltrated with Lowicryl (HM20) resin before polymerising at -50°C.

### Light Microscopy

Brightfield microscopy was carried out on a Zeiss 880 confocal microscope using the TPMT channel and x40 water immersion objective.

### Dot blots

Horseradish peroxidase or *in vitro* generated CAV1-GFP or CAV1-GFP-APEX2 were dotted onto nitrocellulose in a volume of 10 μl to give the indicated protein amounts. After drying, nitrocellulose was incubated with a blocking solution of 5% BSA. After washing with PBS, the nitrocellulose was incubated with the DAB solutions, then with or without the APEX-Gold enhancement process.

### Western blotting of Tpm3.1 APEX2 cells

Triplicates of Tpm3.1-APEX2 +/- and -/- PMEFs were grown on 6 cm dishes until full confluency was reached. Cell lysates were harvested in 4°C RIPA buffer with protease inhibitor (cOmplete, EDTAfree Protease Inhibitor Cocktail, Merck) and homogenised by sonication for 30 sec. Protein concentration was measured with Precision Red Assay (Cytoskeleton). Laemmli sample buffer (Biorad) was added 1:4 (v/v) and lysates were boiled at 95°C for 10 min. Samples were run at 100 V for 90 min on 10% polyacrylamide SDS-PAGE gels in running buffer. Gels were semi-dry transferred in transfer buffer to PVDF membranes pre-activated with 100% methanol. Membranes were blocked in 5% skim milk in TBS for 1 hour and probed with mouse γ/9d 2G10.2 (1:1000, MERCK MABT1335), rabbit anti mouse IgG (1:3000, Abcam ab97046), rabbit α-tubulin (1:3000) (Abcam ab52866) and goat anti rabbit IgG (1:5000) (Biorad 170-6515) antibodies sequentially for 1 hour. Luminata Crescendo Western HRP substrate (Merk) was used for imaging on a Chemicoc MP imaging system (Biorad). Band densitometry was quantified (ImageJ) and normalised to α-tubulin control.

### Cell-free expression and particle characterization, cell incubation

#### Leishmania tarentolae

cell-free lysate was produced, and cell-free protein expression was performed as described by (Hunter et al., 2018). Briefly, *Leishmania tarentolae* Parrot strain was obtained as LEXSY host P10 from Jena Bioscience GmbH, Jena, Germany and cultured in TBGG medium containing 0.2% v/v Penicillin/Streptomycin (Life Technologies) and 0.05% w/v Hemin (MP Biomedical). Cells were harvested by centrifugation at 2500 x g, washed twice by resuspension in 45 mM HEPES, pH 7.6, containing 250 mM Sucrose, 100 mM Potassium Acetate and 3mM Magnesium Acetate and resuspended to 0.25 g cells/g suspension. Cells were placed in a cell disruption vessel (Parr Instruments, USA) and incubated under 7000 KPa nitrogen for 45 min, then lysed by rapid release of pressure. The lysate was clarified by sequential centrifugation at 10 000 x g and 30 000 x g and anti-splice leader oligonucleotide was added to 10 µM. The lysate was then desalted into 45 mM HEPES, pH 7.6, containing, 100 mM Potassium Acetate and 3 mM Magnesium Acetate and snap-frozen until required.

Cell-free lysate was supplemented with a feeding solution containing nucleotides, amino acids, T7 polymerase, HEPES buffer, and a creatine/creatine kinase ATP regeneration system at a ratio of lysate to feed solution of 0.21, and a final Mg^2+^ concentration of 6 mM. Purified plasmid DNA, at a concentration of 1000 ng/µL, was added to the expression reaction at a ratio of 1:9 (v/v), and the reaction allowed to proceed for 3 hour at 27°C. When visualisation of non-GFP-tagged expressed protein was desired, the FluoroTect− GreenLys *in vitro* Translation Labeling System (Promega) was used: labeled tRNA was diluted 1:10 from the supplied material and added to expression reactions at a ratio of 1:9 (v/v). Fluorescently tagged/labeled expressed protein was detected both before and after SDS-PAGE using a Chemidoc MP imaging system (Bio-Rad, Laboratories Pty. Ltd.,Gladesville, NSW, Australia) as described in (Hunter et al., 2018).

### FCS method

Single-molecule fluorescence counting methods were used to compare the oligomeric state of the expressed CAV1-GFP-APEX2 construct with the known state of GFP-labeled CAV1 (GFP-CAV1) Comparison of the brightness values obtained for the two constructs suggests that particles produced using the CAV1-GFP-APEX2 construct comprise approximately 110 GFP-CAV1-APEX2 molecules, approximately 80% of those seen in previous studies with GFP-CAV1

Single molecule spectroscopy was performed using an AttoBright instrument optimised for the detection of GFP (Brown et al, 2019) A 488 nm laser was focussed into the sample solution using a C-Apochromat 40x/1.2 W water-immersion objective lens (Zeiss) and the fluorescence emission was filtered using a 500-550 nm bandpass filter. Samples were diluted 1:5 directly from cell-free expression with Buffer EB and placed in a custom-made silicone 192-well plate with a 70 × 80-mm glass coverslip (ProSciTech). For single-molecule burst brightness analysis, the frequency of events for each range of GFP fluorescence intensity was counted and plotted on a histogram.

Cell-free reaction product was diluted (1:1) with DMEM and incubated on cells for 5 min at 37°C with 5% CO2. Cells were then processed for DAB treatment and APEX-Gold enhancement as stated above.

## STAR Methods Table

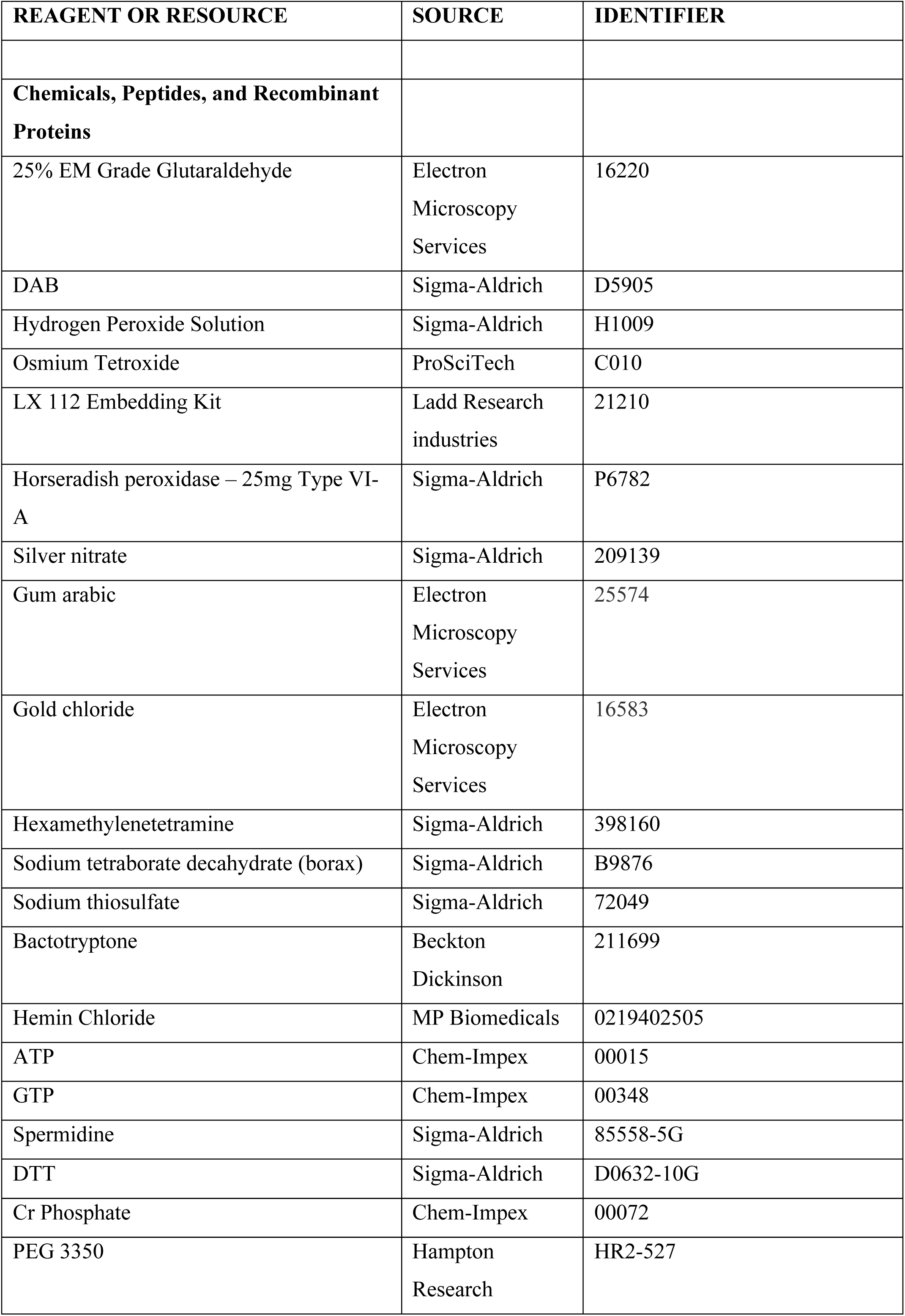
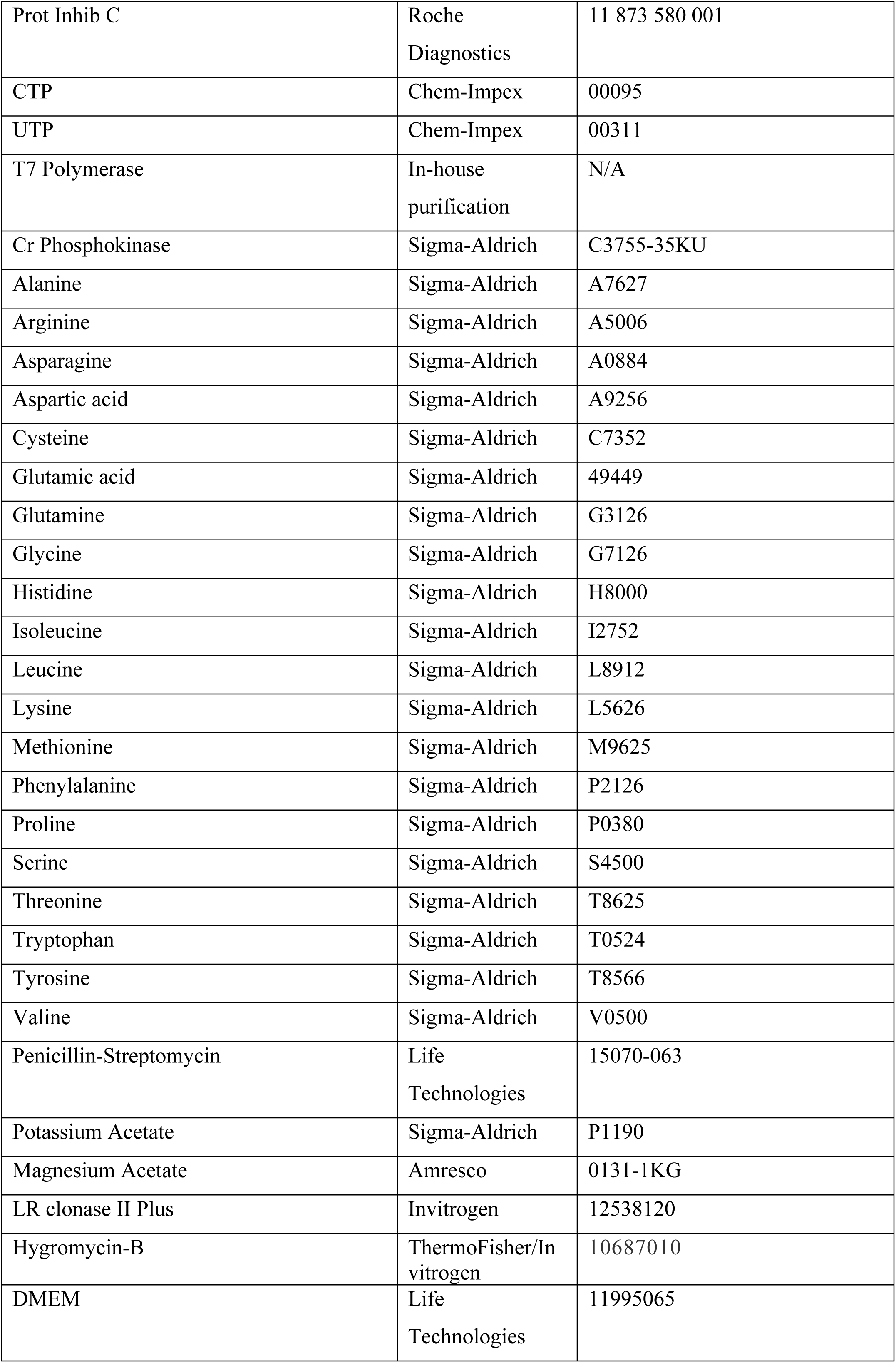
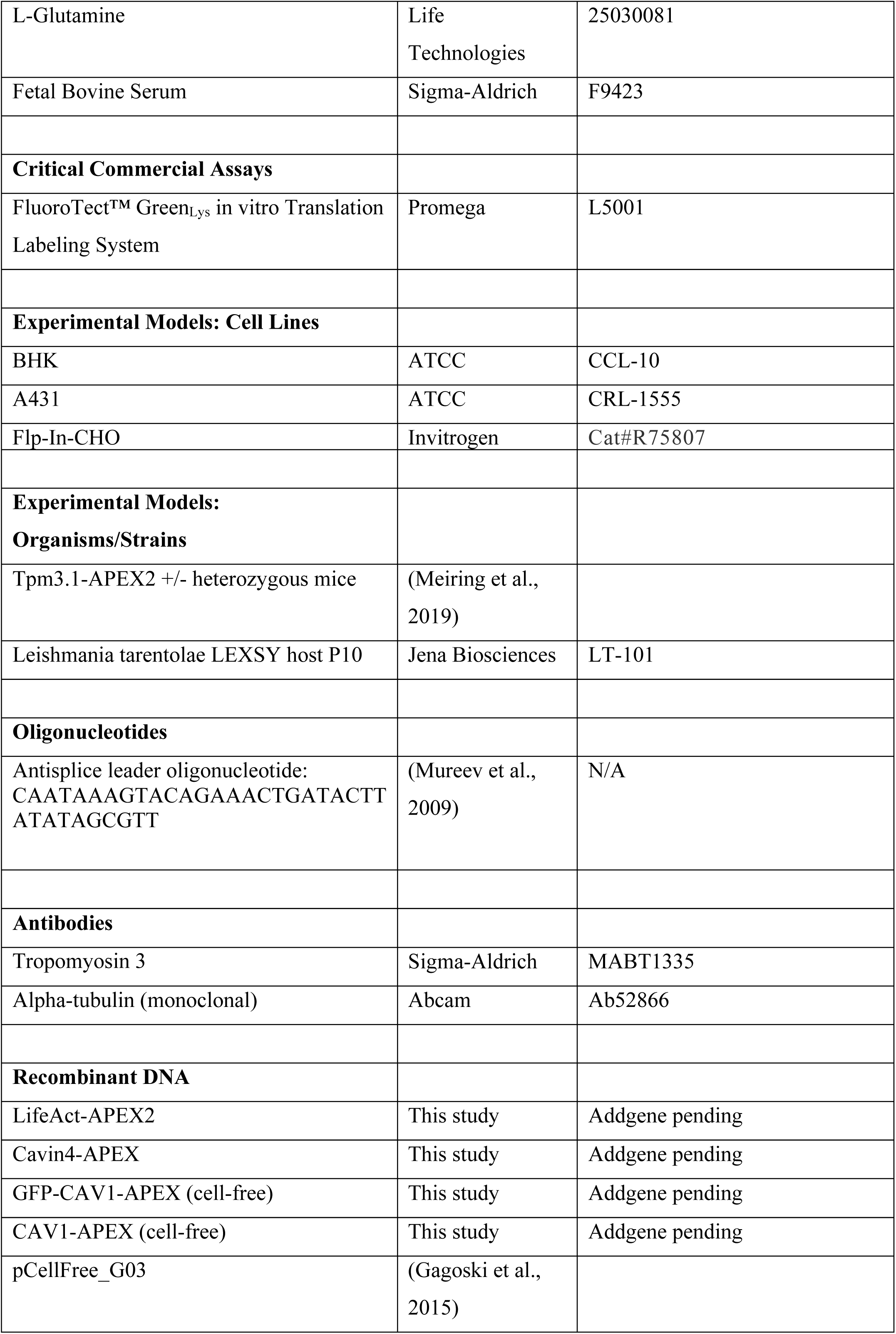
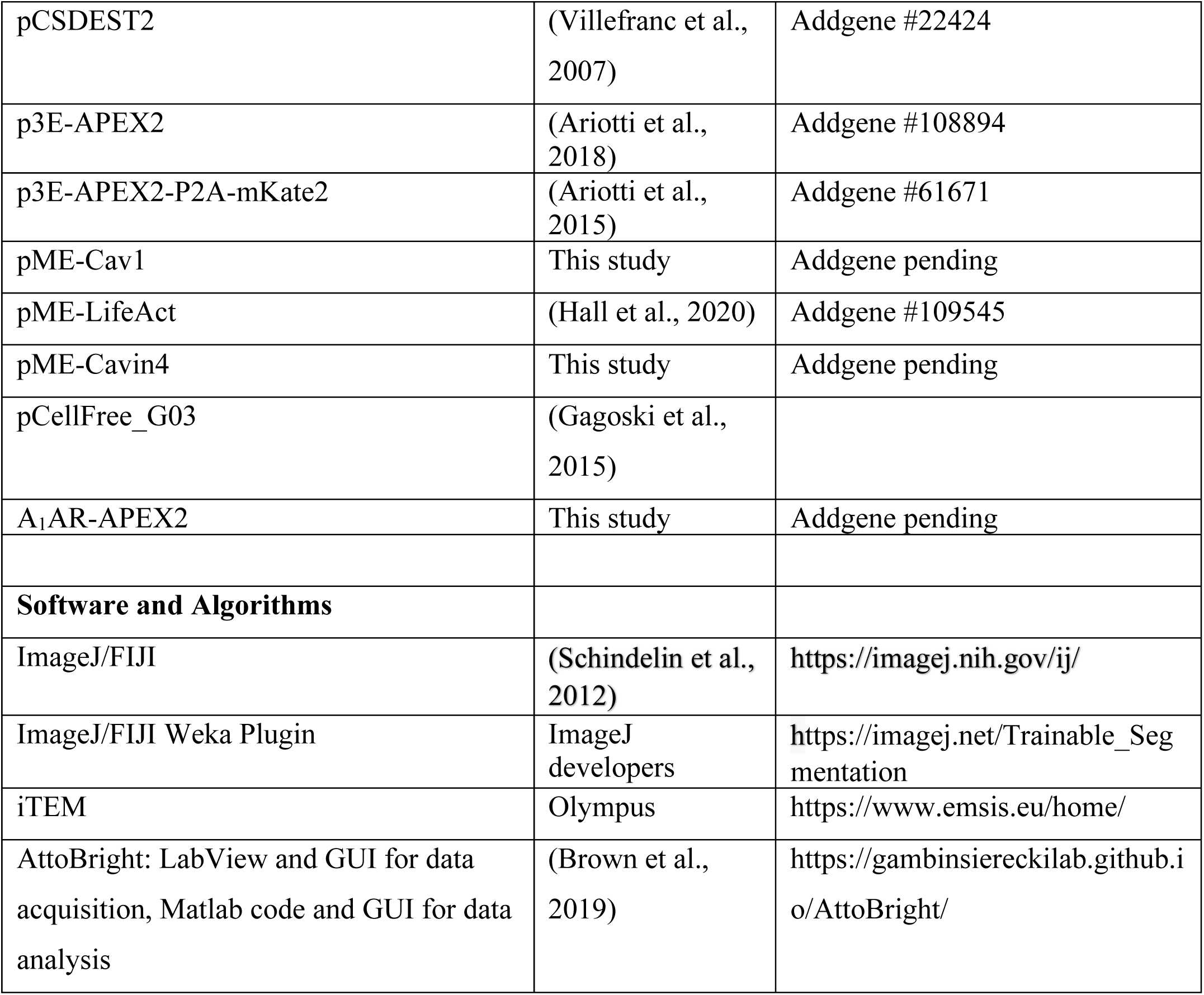

---

**Supplementary Figure 1.**
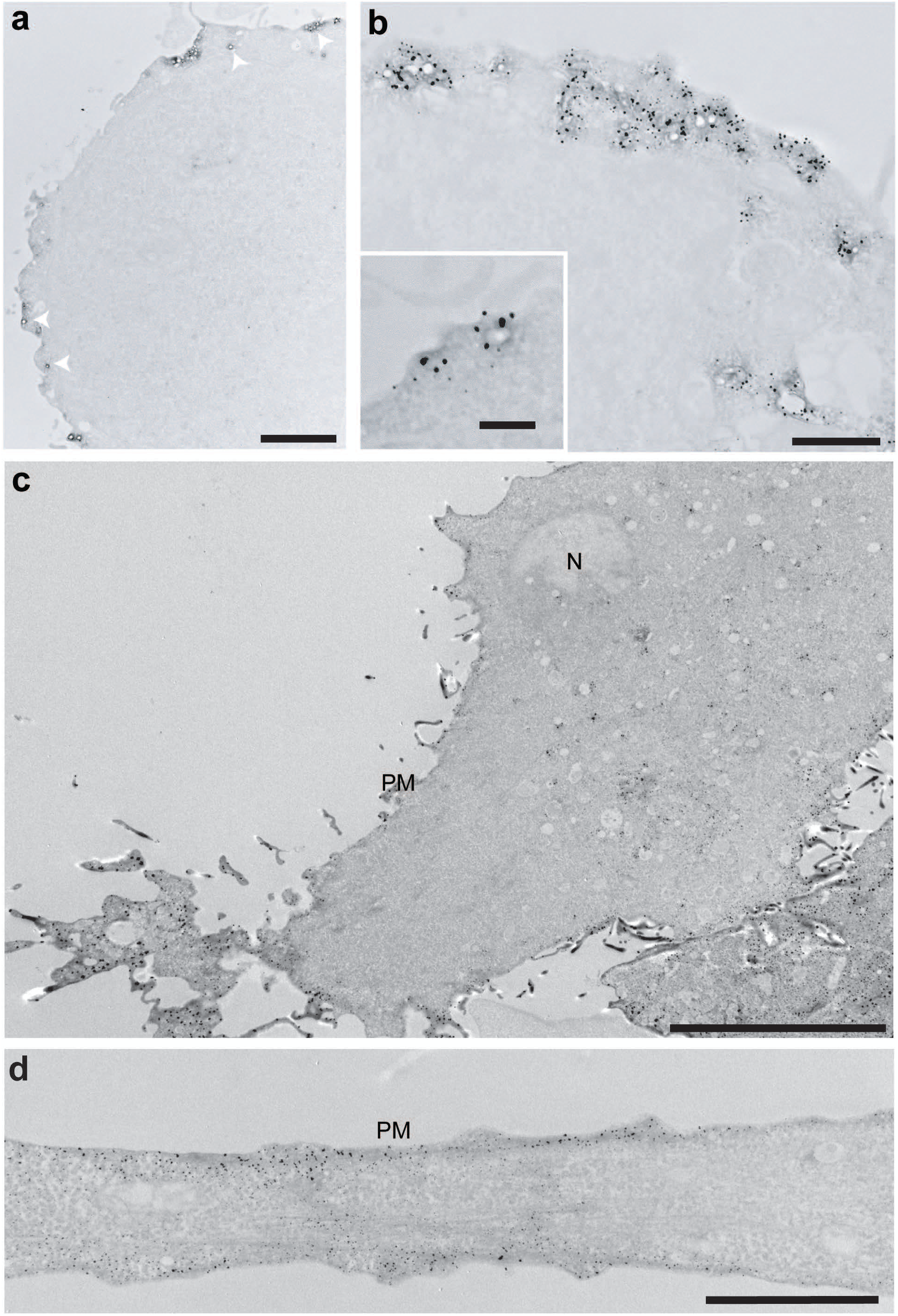
**A,B;** Low and higher magnification views respectively of Cavin4-APEX2 labeling using the APEX-Gold method without gum arabic. Arrows, caveolae labeled with gold. **C,D;** Low and higher magnification examples of labelling of actin filaments with LifeAct-APEX2 with APEX-Gold enhancement. N, nucleus; PM, plasma membrane. Bars, A, 1 µm; B, 1 µm, B inset, 200 nm; C, 10 µm; D, 2 µm.

**Supplementary Figure 2.**
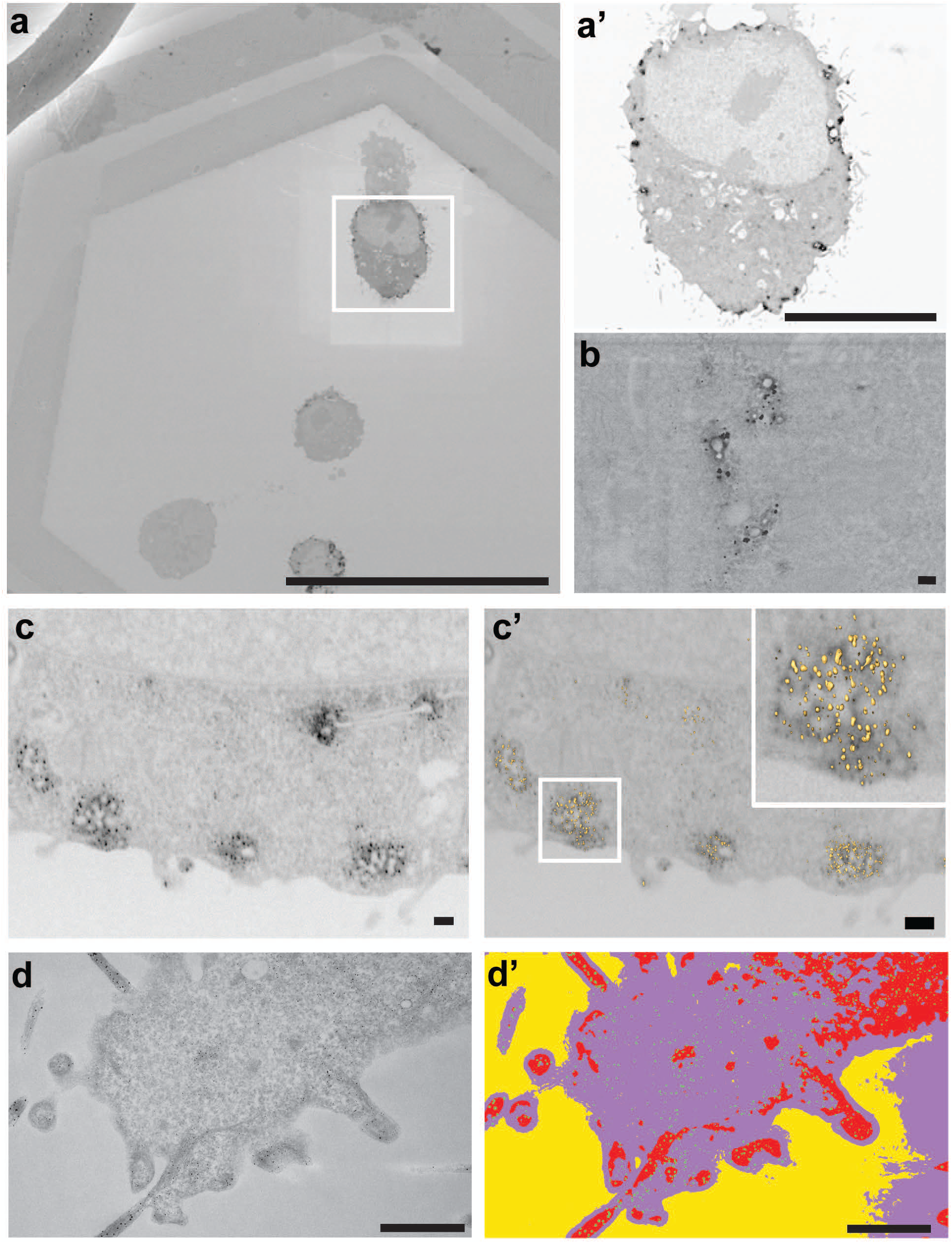
**A-B;** SEM analysis of sections of BHK cells expressing Cavin4-APEX2 (Raith eLine Plus). **C-C’;** FIB analysis of same experiment followed by automated segmentation. **D;** Actin filaments labelled with LifeAct-APEX2 followed by APEX-Gold enhancement. D’; shows segmentation of the DAB product and the gold labeling using FIJI/Weka automated segmentation of the image shown in **D**. Bars,A, 50 µm,A’, 10 µm; B, 100 nm; C,C ‘, 200 nm; D,D ‘, 1 µm.

**Supplementary Figure 3.**
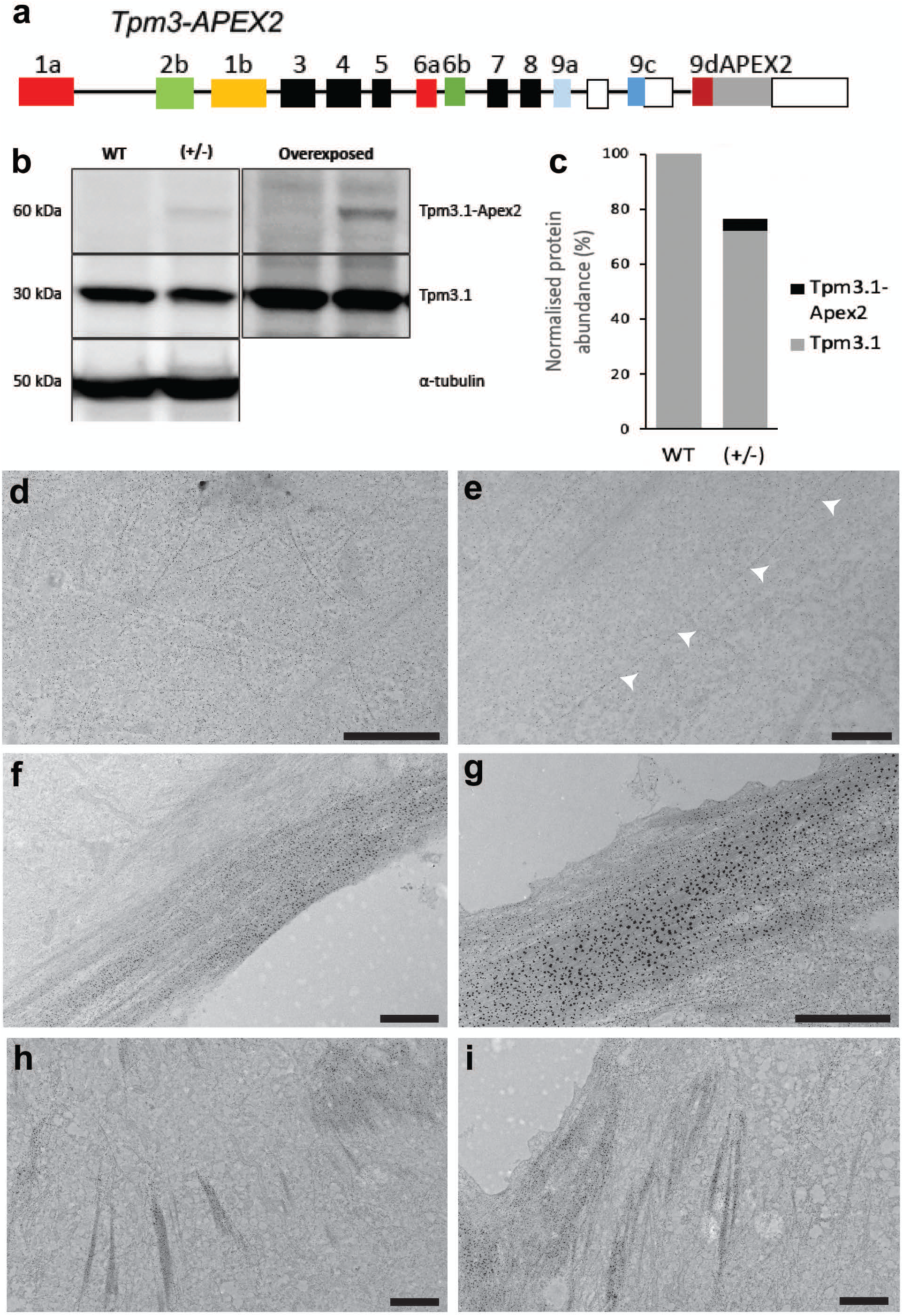
**A;** Genomic organization of the mouse line expressing Tpm3.1 C-terminally tagged with APEX2 (Meiring et al, 2019). B; Western blot detection, and C; quantitation, of endogenous and APEX2-tagged Tpm3.l. D-1; Additional representative micrographs of Tpm3.l-APEX2 labeling with APEX-Gold enhancement. Arrows, particle labeling following individual actin filaments. Bars, A, 2 µm; B, 1 µm; C-F, 2 µm.

**Supplementary Figure 4.**
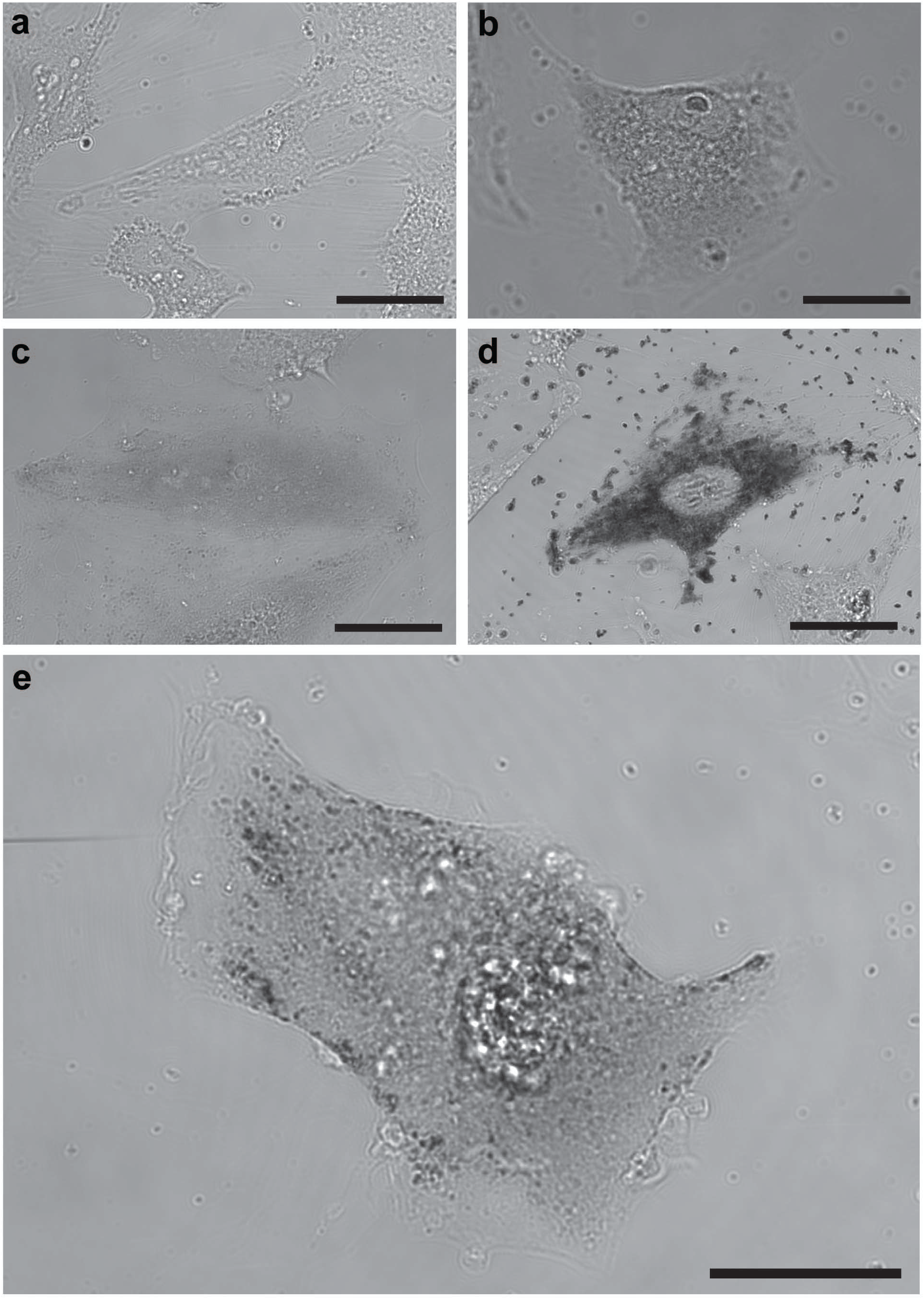
Brightfield images of BHK cells with various treatments. **A;** untransfected BHK cell treated DAB, AG/AU and gum arabic. **B;** BHK cell transfected with Cavin4-APEX2 treated with DAB. C; BHK cell transfected with Cavin4-APEX2 treated with DAB, AG and gum arabic. **D;** BHK cell transfected with Cavin4-APEX2 treated with DAB and AG/AU. **E;** BHK cell transfected with Cavin4-APEX2 treated with DAB, AG/AU and gum Arabic. Bars, A-E, 50 µm.

**Supplementary Figure 5.**
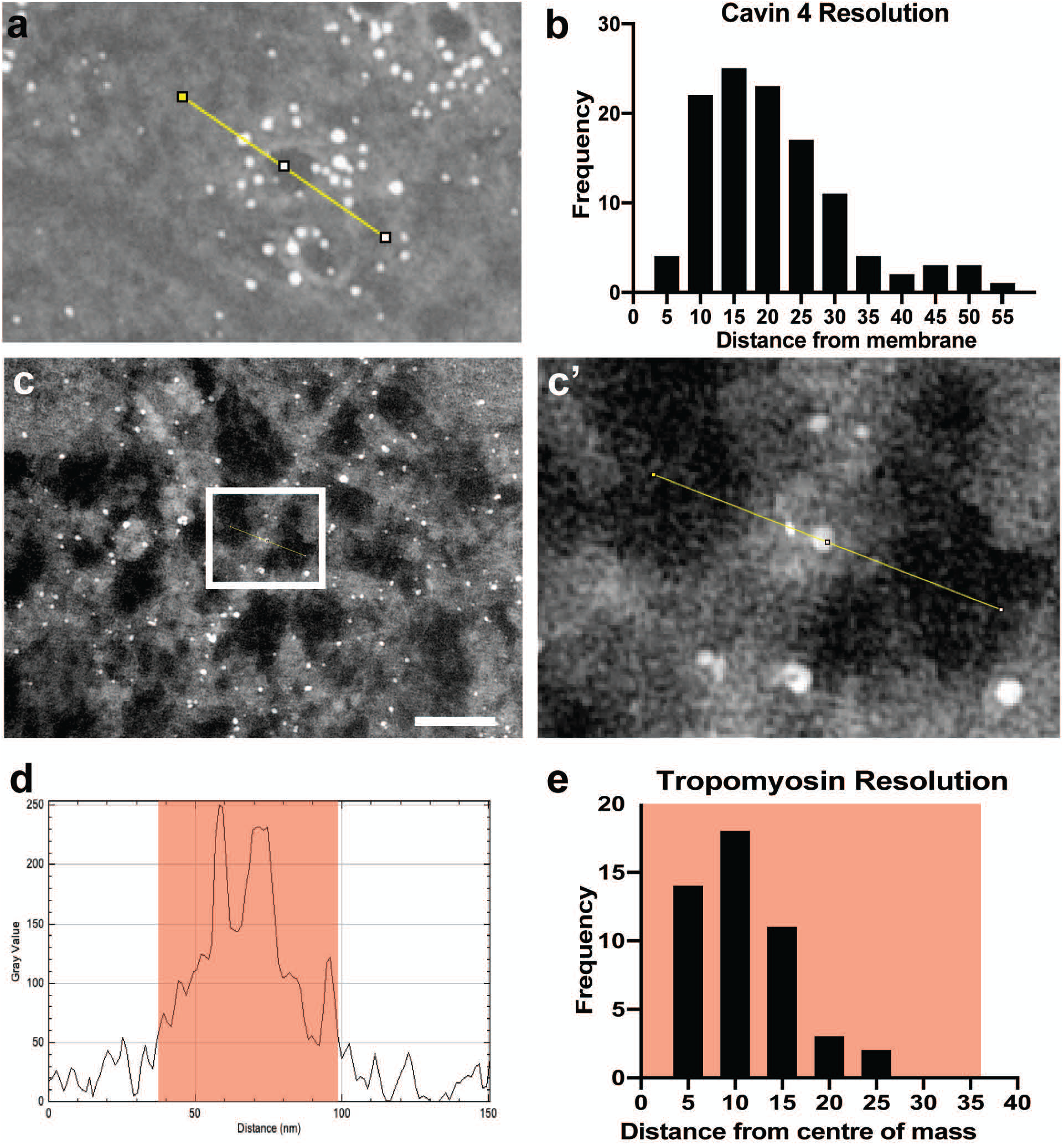
**A;** Micrograph of Cavin4-APEX2 labeling enhanced with APEX-Gold. Linescans (FITI) across caveolae were used to measure the distance from centre of gold particles to the caveolar membrane. **B;** Shows the distribution ofparticle label resolution. **C,C’;** Tpm3.1-APEX2 labeling enhanced with APEX-Gold, also analyzed with linescans to measure distance from particles centre to the actin bundle centre of mass. **D;** Representative output from linescan showing edges of bundle highlighted in orange and the two peaks within indicative ofAPEX-Gold. E; Shows the distri bution of Tpm3.1-APEX2 resolution, and half the average width of bundles highlighted in orange. Bars, C, 50 nm.

**Supplementary Figure 6.**
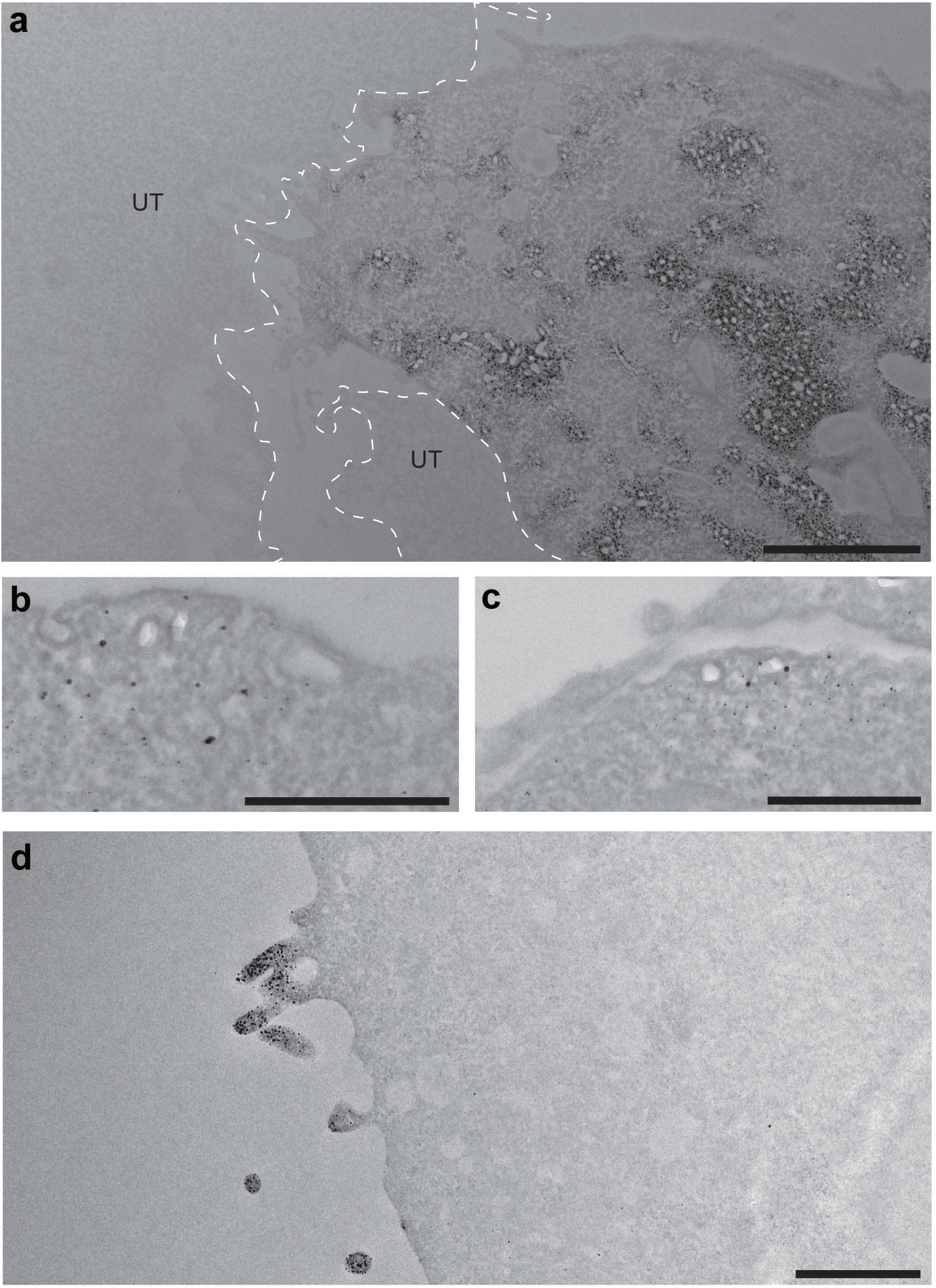
**A;** An example of a cavin4-APEX2 transfected cell immediately adjacent to a untransfected cell (UT, highlighted by dashed line). Note the absence of particulate reaction product within the untransfected cell. **B,C;** Fast frozen and freeze substituted cells embedded in lowicryl showing Cavin4-APEX2 labeling. D; A1AR-APEX2 stably expressing CHO cells, processed with APEX-Gold enhancement without gum arabic showing labelling at specific areas of plasma membrane. UT, untrans fected cell. Bars, A, 2 µm; B-C, 500 nm; D, 1 µm.

**Supplementary Figure 7.**
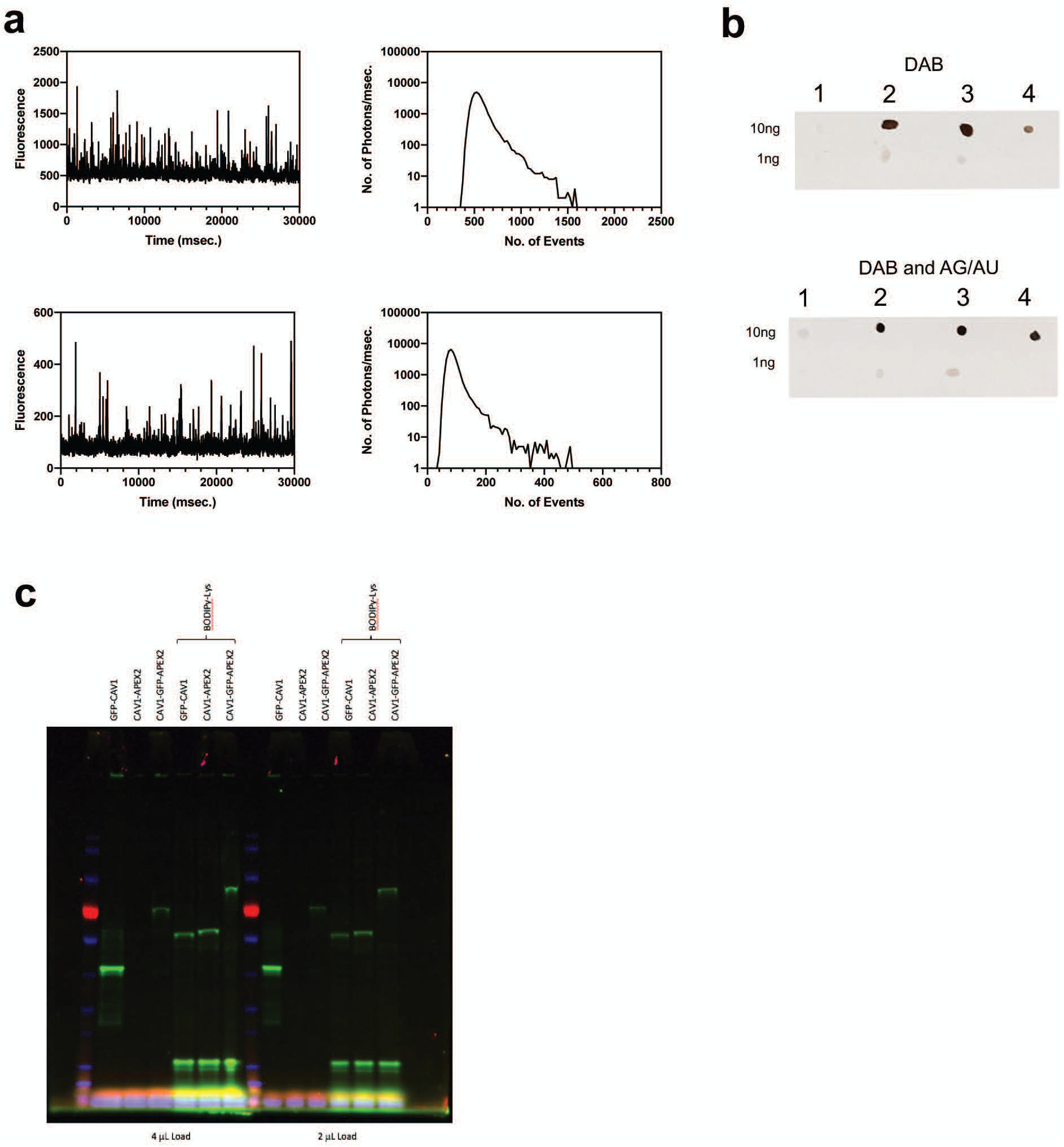
**A;** FCS analysis of cell-free synthesized GFP-CAV1-APEX2. **B;** Dot blots comparing two concentrations ofthe indicated in vitro synthesized proteins (1; GFP-Cavl, 2; Cavl-APEX2, 3; GFP-CAV1-APEX2) or commercial horseradish peroxidase (4; HRP) treated with DAB or DAB and AG/AU. C; Expression of the indicated constructs followed by in-gel fluorescence detection using the GFP fluorescence or with fluorescently labeled lysine (BODIPY-lysine).

## Notes

### Competing Interest Statement

The authors have declared no competing interest.

